# The zinc finger transcription factor Sall1 is required for the early developmental transition of microglia in mouse embryos

**DOI:** 10.1101/2022.02.12.480224

**Authors:** Earl Parker Scott, Emma Breyak, Ryuichi Nishinakamura, Yasushi Nakagawa

## Abstract

Microglia play many critical roles in neural development. Recent single-cell RNA-sequencing studies have found diversity of microglia both across different stages and within the same stage in the developing brain. However, how such diversity is controlled during development is poorly understood. In this study, we first found the expression of the macrophage mannose receptor CD206 in early-stage embryonic microglia on mouse brain sections. This expression showed a sharp decline between E12.5 and E13.5 across the central nervous system. We next tested the roles of the microglia-expressed zinc finger transcription factor SALL1 in this early transition of gene expression. By deleting *Sall1* specifically in microglia, we found that many microglia continued to express CD206 when it is normally downregulated. In addition, the mutant microglia continued to show less ramified morphology in comparison with controls even into postnatal stages. Thus, SALL1 is required for early microglia to transition into a more mature status during development.

## INTRODUCTION

Microglia are resident immune cells in the central nervous system (CNS) derived from Cx3cr1-expressing, erythromyeloid progenitors in the yolk sac (Ginhoux et al., 2010; Gomez Perdiguero et al., 2015; Kierdorf et al., 2013; Schulz et al., 2012). These cells appear in the CNS as early as at embryonic day 9.5 (E9.5) and increase their numbers by proliferation and continuous entry until the blood-brain barrier forms. It is well established that microglia play pivotal homeostatic roles in the adult nervous system and control the synapse number and function in developing brain (Hammond, Robinton, & Stevens, 2018; Prinz, Masuda, Wheeler, & Quintana, 2021). Recent studies also suggest that microglia in embryonic brains play roles in early neural development including neurogenesis and neuronal migration as well as axon growth and guidance (Nandi et al., 2012; Arnò et al., 2014; Squarzoni et al., 2014; Thion & Garel, 2020). Aberrant functions of microglia could lead to a variety of neurodevelopmental disorders including schizophrenia and autism spectrum disorders (Bilbo & Stevens, 2017). In addition, maternal immune activation or inactivation, both of which alter microglia, also disrupts normal brain development (Cunningham, Martinez-Cerdeno, & Noctor, 2013; Tronnes et al., 2016; Bilbo, Block, Bolton, Hanamsagar, & Tran, 2018; Choi et al., 2016). However, the underlying molecular and cellular mechanisms by which early microglia influence neural cells are still poorly understood, and there are some discrepancies between the results of published studies (Cunningham et al., 2013; Squarzoni et al., 2014; Arnò et al., 2014; Nandi et al., 2012). Recent single-cell RNA sequencing analyses have indicated step-wise changes in gene expression during embryonic and postnatal development of microglia (Li et al., 2019; Matcovitch-Natan et al., 2016; Thion et al., 2018; Masuda et al., 2019; Masuda, Sankowski, Staszewski, & Prinz, 2020). In addition, subpopulations of microglia exist even at the same developmental stage (Hammond et al., 2019). Therefore, to better understand the roles of microglia in individual neurodevelopmental events, it is essential to determine how microglia development is regulated both spatially and temporally. In this study, we first characterized the temporal patterns of microglia gene expression by focusing on the macrophage mannose receptor, CD206 (encoded by the *Mrc1* gene). *CD206/Mrc1* has been considered to be a marker for non-microglial, brain border-associated macrophages (BAMs) during embryonic development (Utz et al., 2020), although a bulk RNA-sequencing found that *Mrc1* is enriched in early embryonic stage (“early microglia”) until E12.5 (Matcovitch-Natan et al., 2016). In adult microglia, expression of CD206 is reduced in the absence of the fractalkine receptor Cx3cr1 (Piirainen et al., 2021), and is induced by inhibition of TGFβ signaling (von Ehr et al., 2020). CD206 is also induced in spinal cord injury (Kisucká, Bimbová, Bačová, Gálik, & Lukáčová, 2021), stroke (Hu et al., 2012) and under chronic restraint stress (Piirainen et al., 2021), and may have implications in synaptic pruning (Piirainen et al., 2021). However, spatial and temporal distribution of CD206 in developing brain and its role in neural development have not been explored. By immunohistochemistry, we found that a majority of Iba1-positive, parenchymal immune cells in early embryonic CNS express CD206. The percentage of CD206-expressing Iba1+ cells dropped dramatically during embryogenesis between E12.5 and E13.5. Coincidentally, the transcription factor *Sall1*, a signature microglial gene, is upregulated at a similar timing (Matcovitch-Natan et al., 2016; Thion et al., 2018). *Sall1* is a mammalian homologue of the Drosophila homeotic gene *spalt* and encodes a zinc-finger transcription factor that plays crucial roles in the development of many organs including the kidney (Nishinakamura et al., 2001). Heterozygous mutations in the human *SALL1* gene cause Townes-Brock Syndrome (Townes & Brocks, 1972; Kohlhase, Wischermann, Reichenbach, Froster, & Engel, 1998), which involves multiple organ systems and often causes intellectual disability (Bardakjian, Schneider, Ng, Johnston, & Biesecker, 2009; Powell & Michaelis, 1999). We validated SALL1 expression on embryonic brain sections by immunohistochemistry, and tested if *Sall1* is required for the early transition of microglia status by deleting the gene specifically in microglia. We found that without *Sall1*, many microglia continued to express CD206 into postnatal stages. Morphology of microglia also became less ramified compared with controls even into postnatal stages. These results show that SALL1 is a critical regulator of normal progression of microglia development early in embryonic brains, which may have implications on the microglia’s roles in specific aspects of neural development.

## RESULTS

### Temporal pattern of CD206 expression in parenchymal Iba1+ cells in embryonic CNS

To determine the spatial and temporal pattern of microglia development, we analyzed the expression of CD206 (encoded by the *Mrc1* gene), which is expressed in *Cx3cr1*-expressing, brain immune cells at E10.5 and E12.5 but is downregulated by E13.5 in bulk RNA-sequencing (Matcovitch-Natan et al., 2016). With immunohistochemistry, we found that at E10.5 and E11.5, 60-70% of parenchymal Iba1-positive cells also expressed CD206 (Fig.1A-F). At E12.5, similar percentages of Iba1+ cells in the parenchyma of the neocortex, retina, thalamus and ganglionic eminences still expressed CD206 (Fig.1G-N) along with the microglia-specific marker, P2RY12 (Fig.1O). By E13.5, however, few Iba1+ cells in the CNS parenchyma expressed CD206 in all of the above CNS regions (Fig.1P,Q), which was consistent with the RNA sequencing data (Matcovitch-Natan et al., 2016). Thus, loss of CD206 expression could be used as a marker for early developmental transition of microglia.

**Figure 1.**
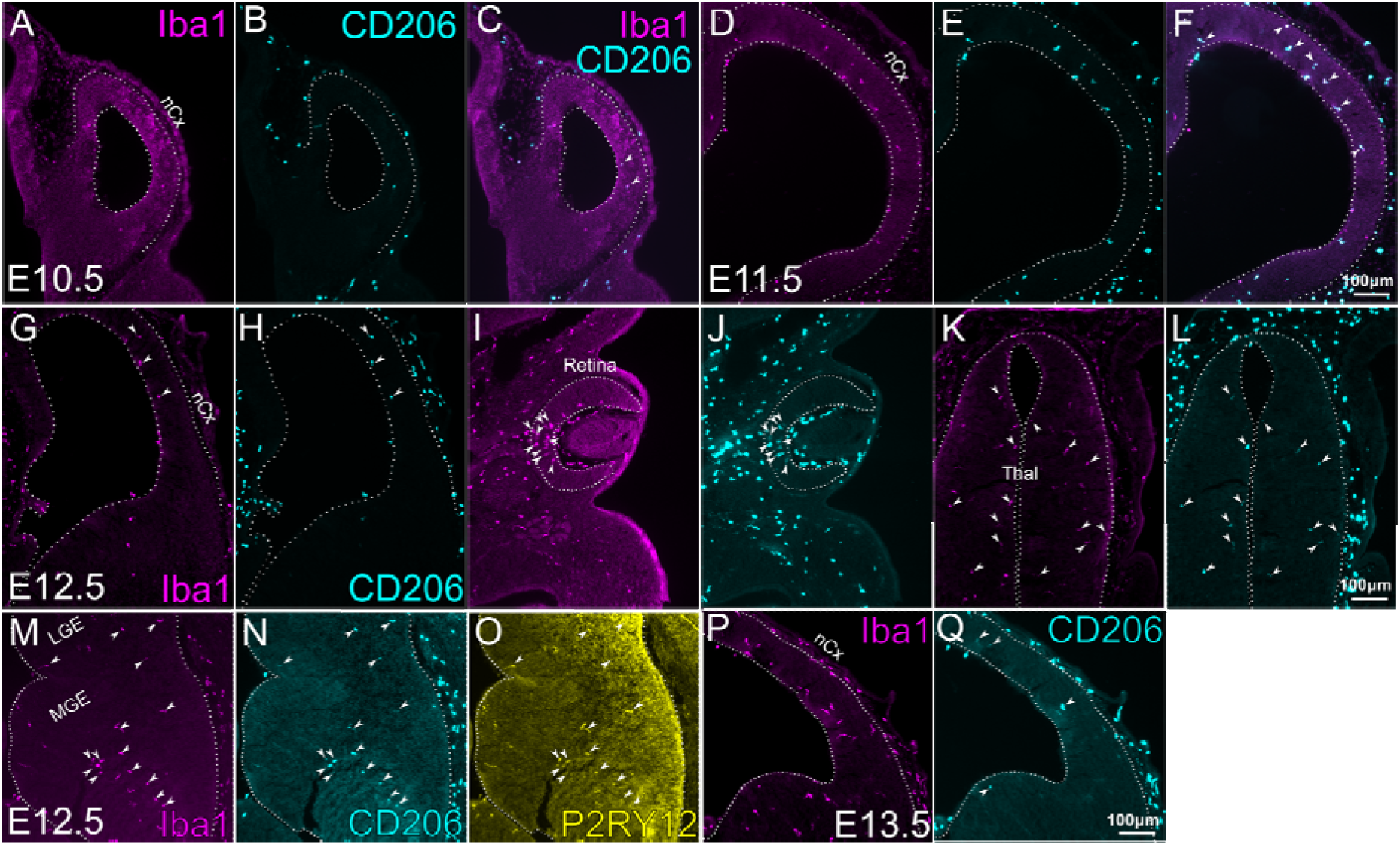
Expression of CD206 in early CNS parenchyma. **A-L.** Double immunostaining of Iba1 and CD206 on frontal sections of mouse embryos at E10.5 (A-C), E11.5 (D-F), and E12.5 (G-L). Various CNS regions are shown for E12.5. G,H; telencephalon, I,J; retina, K,L; thalamus. All images show the right side of the section. Arrowheads indicate cells that express both Iba1 and CD206. **M-O.** Triple immunostaining of Iba1 (M) CD206 (N) and P2RY12 (O) at E12.5 within ganglionic eminences. Arrowheads indicate cells that express all three markers. **P-Q.** Double immunostaining of Iba1 and CD206 on frontal sections of mouse embryos at E13.5. Unlike at E10.5, E11.5 and E12.5, E13.5 CNS has few cells that express both Iba1 and CD206 (arrowheads), despite an increased density of Iba1+ cells. nCx, neocortex; MGE, medial ganglionic eminence; LGE, lateral ganglionic eminence. Scale bars, 100μm.

### Transcription factor SALL1 is expressed in embryonic microglia as early as at E12.5

In order to determine the intrinsic mechanisms regulating microglia development in early embryonic stages, we next examined the expression of SALL1, a zinc finger transcription factor required for the maintenance of homeostatic microglia phenotypes in adult cerebral cortex (Buttgereit et al., 2016). Previous RNA-sequencing studies showed that *Sall1* is expressed in microglia and not in BAMs (Buttgereit et al., 2016; Kim et al., 2021), and is strongly unregulated during embryogenesis (Thion et al., 2018; Matcovitch-Natan et al., 2016). In PU.1+ immune cells in the CNS, SALL1 was initially detected by immunohistochemistry at E12.5 in all regions of the CNS that we analyzed, including the neocortex, ganglionic eminences, retina and the thalamus (Fig.2). In addition to PU.1+ immune cells, SALL1 was robustly expressed in radial glia within the ventricular zone of most CNS regions including the cerebral cortex (Harrison, Nishinakamura, Jones, & Monaghan, 2012), ganglionic eminences and spinal cord, but not in the retina and the thalamus (Fig.2). Based on these results, we next sought to determine if SALL1 plays a role in microglia development by conditionally deleting the gene at the earliest stages of its expression without affecting SALL1 expression in neural progenitor cells.

**Figure 2.**
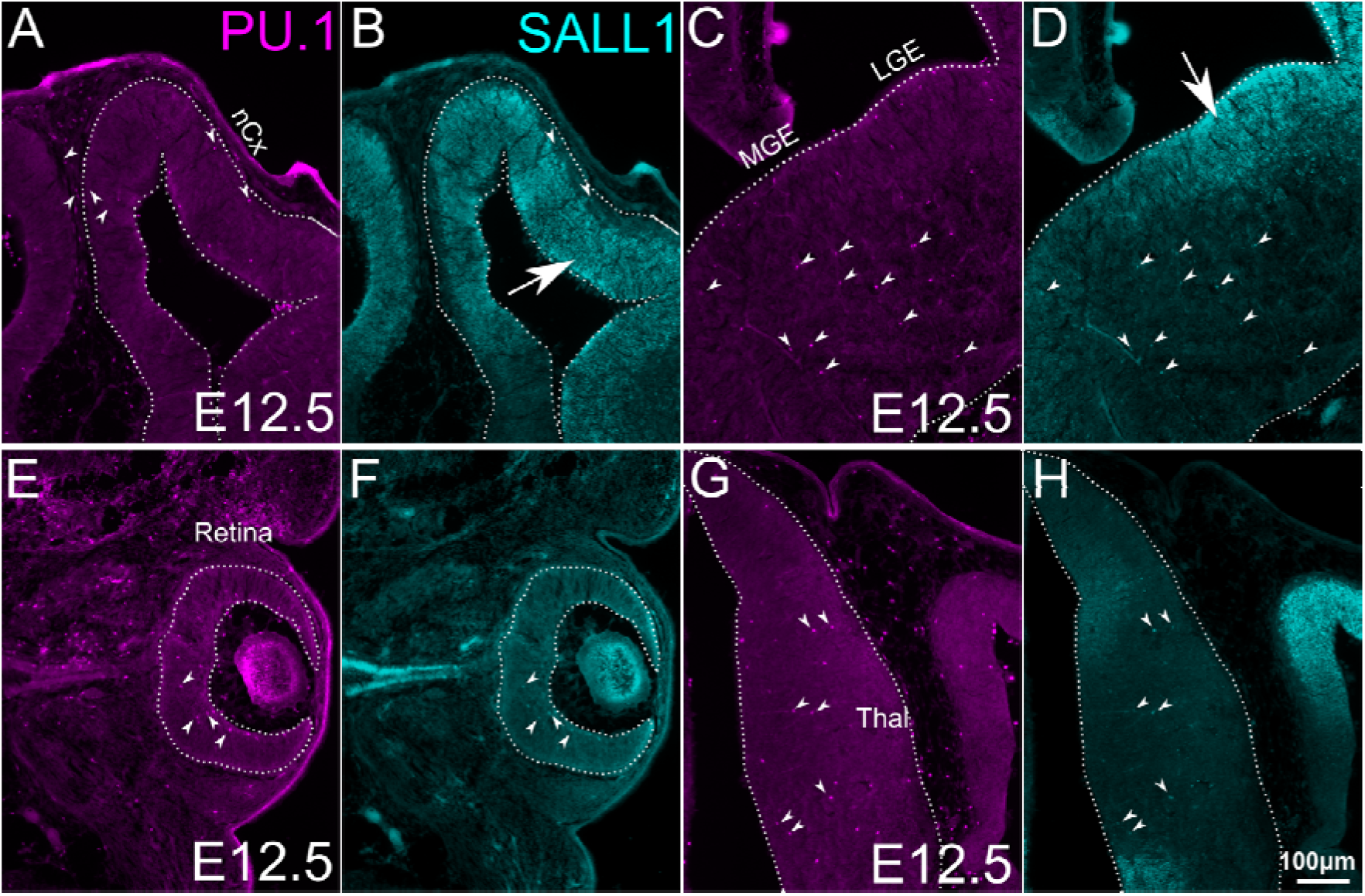
Expression of SALL1 in embryonic CNS. Double immunostaining of SALL1 and PU.1 on frontal sections of mouse embryos at E12.5. Right sides of the brain are shown. A,C,E,G are for PU.1 and B,D,F, H are for SALL1. Arrowheads indicate cells that express all three markers. **A,B.** neocortex (nCx). **C,D.** medial (MGE) and lateral (LGE) ganglionic eminences. **E,F.** retina. **G,H.** thalamus. SALL1 is expressed not only in PU.1-positive immune cells in the CNS, but also in radial glial cells in the neocortex (arrow in B) and LGE (arrow in D), but not in the retina or in the thalamus. Scale bars, 100μm.

### The *Lyve1^Cre^* driver causes recombination in microglia and brain-border associated macrophages

As a constitutive Cre driver for deleting *Sall1* in microglia, we used *Lyve1^Cre^* mice (Pham et al., 2010). Although LYVE1 has been described as a marker for BAMs and lymphatic endothelial cells, and is not expressed in microglia (Jackson, 2003; Pham et al., 2010; Utz et al., 2020), the *Lyve1^Cre^* driver allows tracing the lineage of erythromyeloid progenitor cells in the yolk sac (Lee et al., 2016). Because erythromyeloid cells are the major source of microglia and BAMs (Ginhoux et al., 2010; Gomez Perdiguero et al., 2015; Kierdorf et al., 2013; Schulz et al., 2012), we predicted that the *Lyve1^Cre^* mice might be able to drive the recombination in microglia and BAMs from the earliest stage of their development in CNS. In fact, when *Lyve1^Cre^* mice were bred with a tdTomato reporter mice (Ai14), we found that the reporter was expressed in most Iba1+ cells in CNS parenchyma as early as at E10.5 (Fig.3A,B) and at later stages (Fig.3C,D for E12.5). At E16.5, approximately 90% of Iba1+ cells in the parenchyma of the neocortex expressed tdTomato (Fig.3G,H). The few recombined cells that did not express Iba1 exhibited a tube-like morphology compatible with their lymphatic endothelial identity (yellow arrowheads in Fig.3G,H). Based on this pattern of recombination, we tested if the *Lyve1^Cre^* driver conditionally deletes *Sall1* in microglia without affecting neural cells. Indeed, already at E12.5, *Lyve1^Cre/+^; Sall1^flox/flox^* lacked the expression of SALL1 in PU.1+ parenchymal cells, whereas the expression in radial glial cells in the ventricular zone appeared intact (Fig.3I-L). We detected very few endogenous LYVE1 in parenchymal Iba1+ cells (Fig.3E,F). These results collectively suggest that *Lyve1* is expressed temporally in precursor cells of microglia and BAMs before their entry into the CNS.

**Figure 3.**
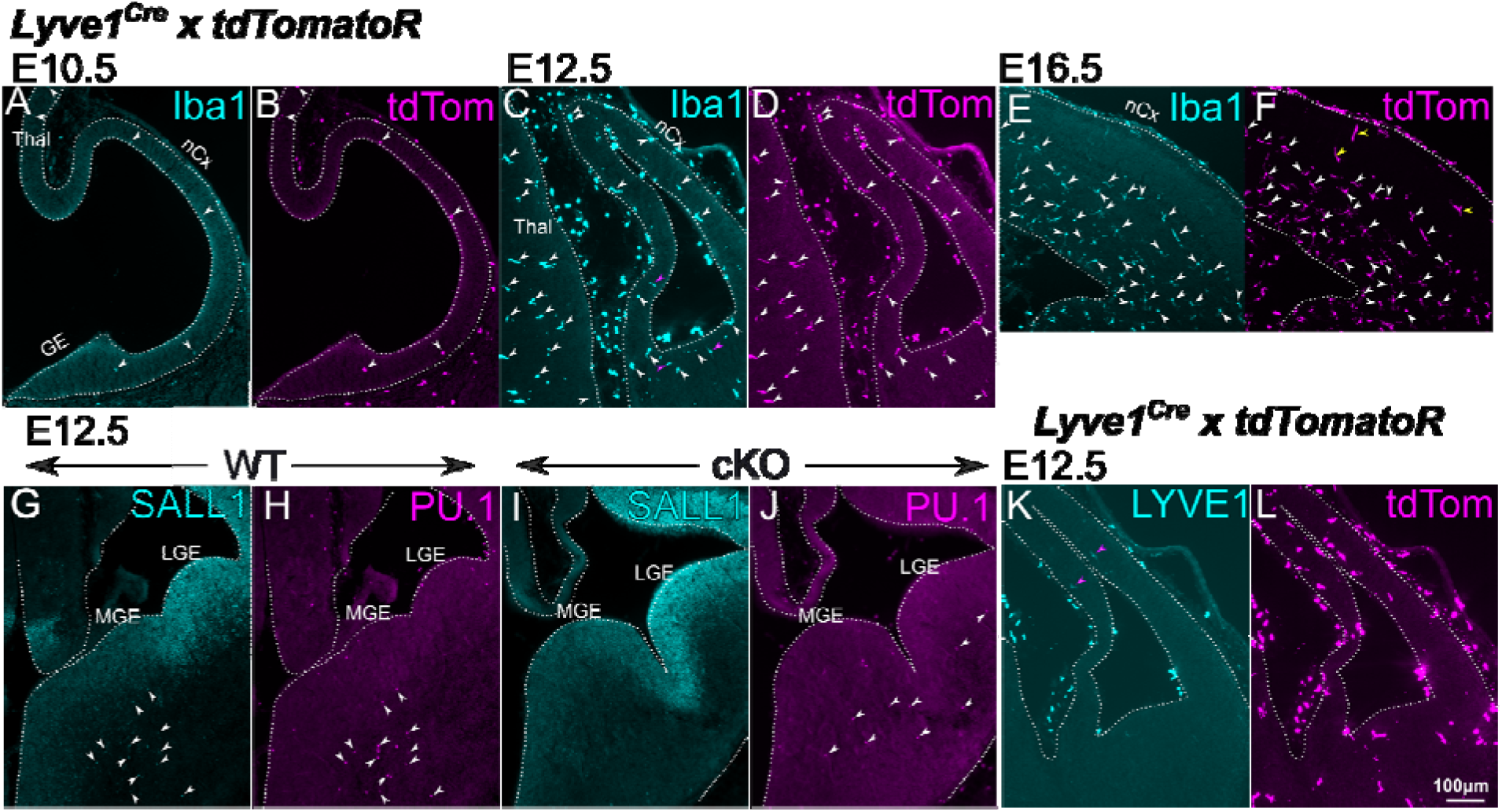
Recombination by the *Lyve1^Cre^* driver. **A-F.** Double immunostaining of Iba1 and the tdTomato in *Lyve1*^*Cre/*+^; tdTomato reporter mice at E10.5 (A,B), E12.5 (C,D) and E16.5 (E,F). Recombination in CNS parenchyma starts to be detected at E10.5 as Iba1-tdTomato double-positive cells (arrowheads). Such cells dramatically increase at E12.5 (C,D). At E16.5, most Iba1+ cells still express tdTomato (white arrowheads) with few cells that express tdTomato but not Iba1 (yellow arrowheads in H). **G-J.** Conditional deletion of *Sall1* using the *Lyve1^Cre^* driver. In wild type brains, SALL1 is expressed in PU.1+ immune cells in the brain parenchyma (arrowheads in G,H), but not in *Sall1* conditional knockout mice. PU.1+ cells (arrowheads in J) do not express SALL1 (I). **K,L.** Very few cells express LYVE1 in the brain parenchyma in the neocortex at E12.5 (arrowheads in K). nCx, neocortex; MGE, medial ganglionic eminence; LGE, lateral ganglionic eminence. Scale bars, 100μm.

### Deletion of *Sall1* results in the sustained expression of CD206

In order to determine if SALL1 plays a role in the developmental transition of early microglia during embryogenesis (Matcovitch-Natan et al., 2016), we next detected CNS cells that expressed both Iba1 and CD206 at E12.5 and E13.5, and compared the ratio of the doublepositive cells among Iba1+ parenchymal cells (Iba1+CD206+/Iba1+) between *Sall1* conditional knockout (cKO) and wild type control mice. At E12.5, 50-75% of parenchymal Iba1+ cells also expressed CD206 in the cortex, lateral and medial ganglionic eminences, thalamus, and the retina (Fig.1G-O). At E13.5, the cortex and ganglionic eminences in wild type animals underwent a dramatic reduction of this ratio to 20-25% (Fig1P,Q, Fig4A,B, I). In the retina, less than 10% still expressed CD206 in E13.5 wild type controls (Fig4E,F,I). In *Sall1* cKO brains, much higher percentages of Iba1+ cells expressed CD206 approximately compared with wild type brains in all regions except the retina (Fig.4C,D,I). In the retina, a much lower percentage (~15%) of Iba1+ cells also expressed CD206 in the cKO mice, but that was still significantly higher than in wild type controls (Fig.4G-I). The sustained CD206 expression continued until postnatal stages (Fig.4K-N). A vast majority of CD206+ cells also expressed the microglia-specific markers, P2RY12 at E16.5 (Fig.5A-F) as well as Tmem119 at P14(Fig.5G.J), and the cKO brain parenchyma still lacked the expression of the BAM marker, LYVE1 (Fig.4I,J) (Utz et al., 2020). These results collectively indicate that mutant Iba1+ cells at least partially retained the identity as microglia and expressed an early microglia marker, CD206. The density of Iba1+ cells was comparable between cKO and wild type brains at both E12.5 and E13.5, whereas the density of CD206+ cells in cKO brains did not drop as much as in wild type between E12.5 and E13.5 (Fig.4J). This suggests that the increased Iba1+CD206+/Iba1+ ratio in *Sall1* cKO brains is due to the persistent expression of CD206 in Iba1+ cells, and not the lack of increase in the number of Iba+CD206-cells. Similar results were obtained when *Cx3cr1^CreER^* allele was used as a driver and CreER was activated by tamoxifen at E11.5 (Fig.S1A-D), validating our results with the *Lyve1^Cre^* driver.

**Figure 4.**
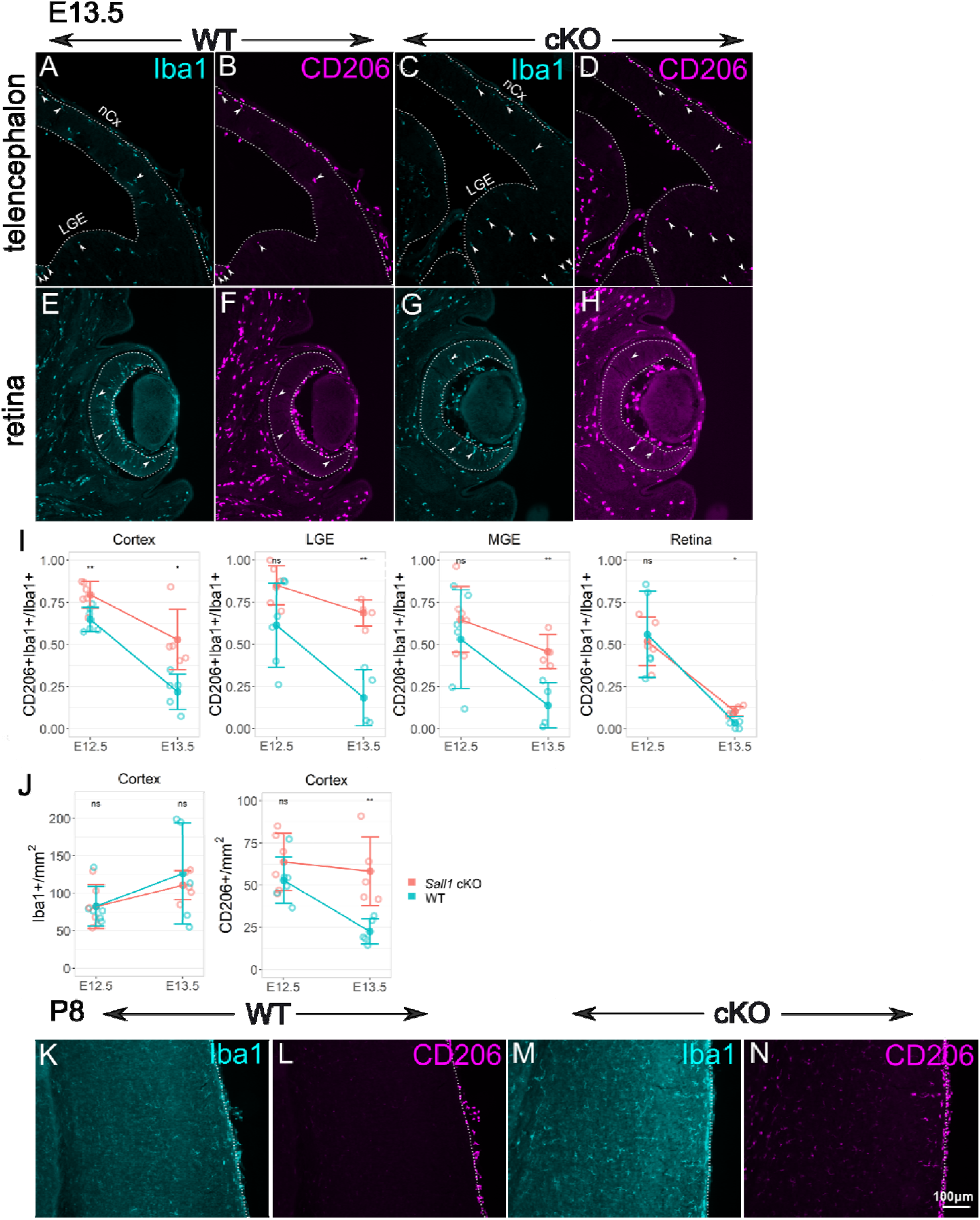
Sustained expression of CD206 in Sall1-deficient microglia. **A-H.** Double immunostaining of Iba1 and CD206 in wild type cortex (A,B) and retina (E,F), and *Sall1* knockout cortex (C,D) and retina (G,H) at E13.5. In wild type controls, few Iba1+ parenchymal cells express CD206, but *Sall1* mutant brains have more CD206+; PU.1+ cells (arrowheads). **I**. Quantitative analysis of the ratio of CD206+Iba1+/Iba1+ cells comparing wild type (green) and *Sall1* mutant (pink) mice in different CNS regions at both E12.5 and E13.5. **, p<0.01; *, p<0.05. **J.** Quantitative analysis of the density of Iba1+ cells and CD206+ cells comparing wild type (green) and *Sall1* mutant (pink) mice in the cortex at both E12.5 and E13.5. **, p<0.01. **K-N.** Double immunostaining of Iba1 and CD206 in wild type (K,L) and *Sall1* knockout (M,N) cortex at P8. Scale bars, 100μm.

**Figure 5.**
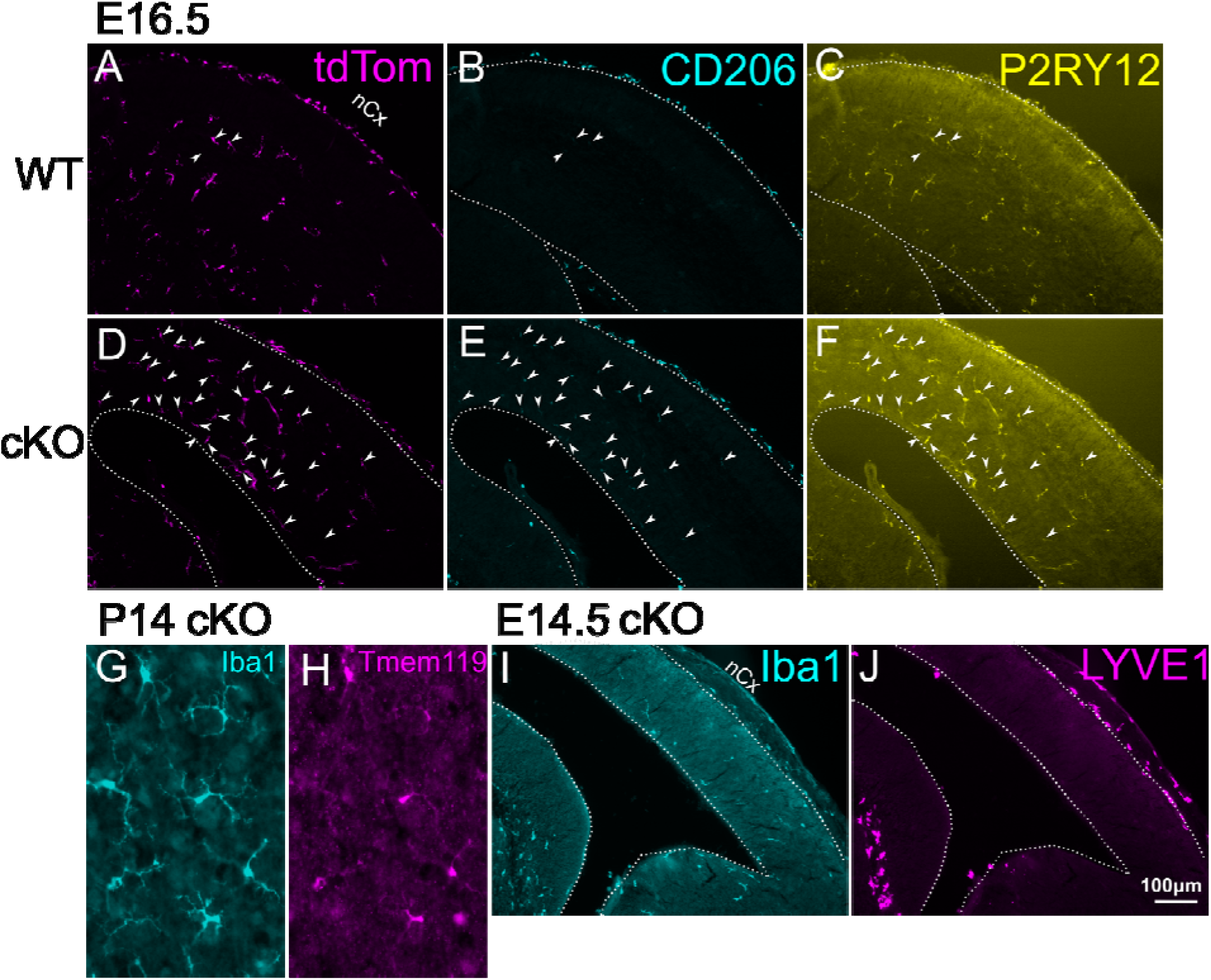
Analysis of marker expression in Iba1+ cells in *Sall1* mutant brains. **A-F.** Triple immunostaining of tdTomato (A,D) CD206 (B,E) and P2RY12 (C,F) in wild type (*Lyve1*^*Cre/*+^; tdTomato reporter; *Sall1^wt/wt^*) and Sall1 cKO (*Lyve1*^*Cre/*+^; tdTomato reporter; *Sall1^flox/flox^*) embryos at E16.5. Arrowheads indicate cells that express all three markers. Most of the persistent CD206+ cells in the parenchyma of the cKO brains are tdTomato+ and also express the microglia marker, P2RY12. **G,H.** Double immunostaining of Iba1 and Tmem119 in *Sall1* knockout cortex at P14, showing that parenchymal cells that express Iba1 still express the microglia marker, Tmem119. **I,J.** Double immunostaining of Iba1 (I) and LYVE1 (J) in *Sall1* cKO brain at E14.5 showing that mutant microglia do not express the BAM marker, LYVE1. Scale bars, 100μm.

### Some axon tracts contain CD206+ microglia even in wild type brains

Although there was a dramatic reduction in the number of CD206+ microglia in wild type CNS between E12.5 and E13.5, some regions near axon tracts persistently expressed CD206. One such region was along the external capsule of the embryonic brain (Fig.6A,B). These cells appeared to be still present in *Sall1* cKO brains (Fig.6C,D). In addition, the corpus callosum in early postnatal wild type brains contained a number of CD206-expressing microglia (Fig.6E,F). These cells still remained in *Sall1* mutants, but they appeared more scattered than in wild type brains (Fig.6G,H).

**Figure 6.**
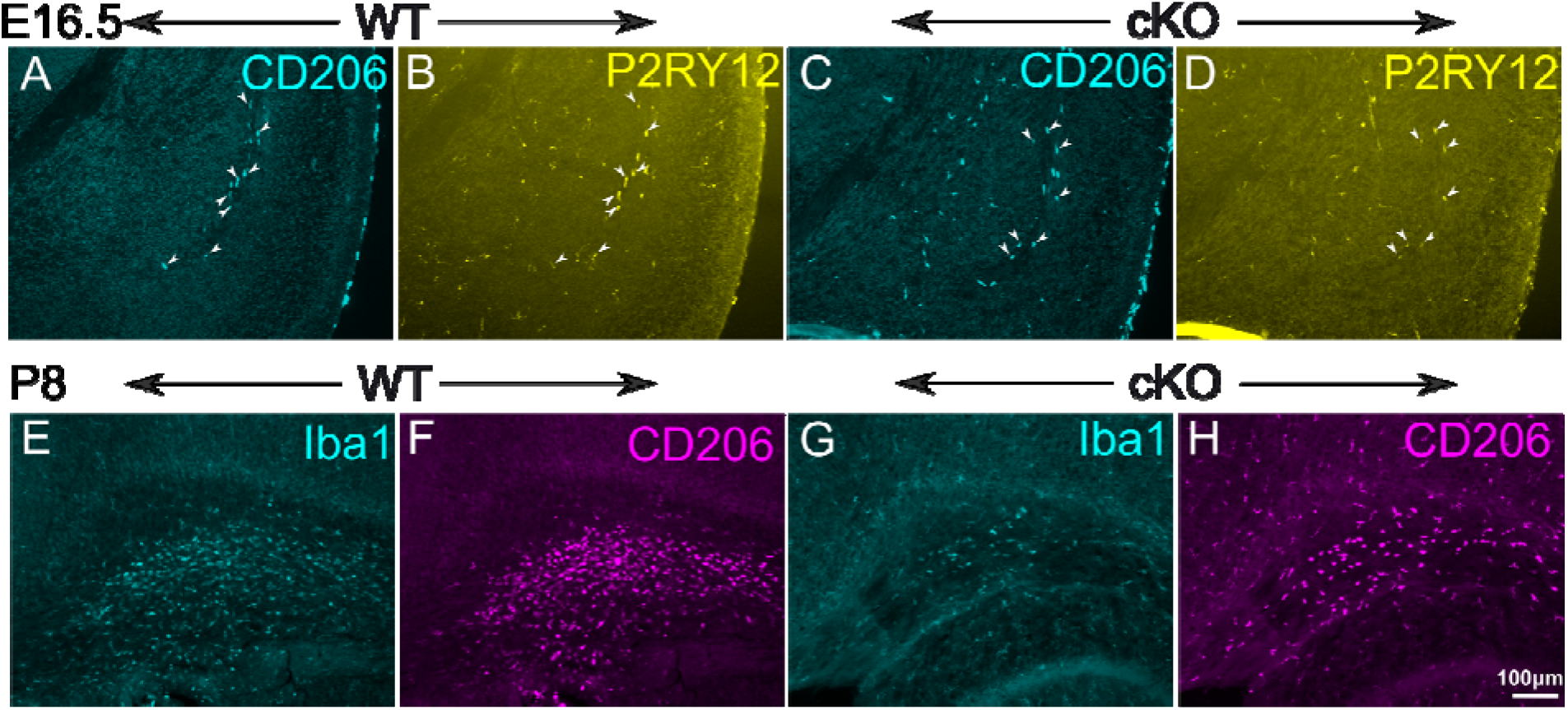
Tract-associated microglia express CD206 in wild type as well as in *Sall1* mutant brains. **A-D.** Double immunostaining of CD206 and P2RY12 in the subpallium of wild type (A,B) and *Sall1* knockout (C,D) mice at E16.5. Arrowheads indicate cells that express both CD206 and P2RY12. Near the external capsule, CD206 is expressed in P2RY12+ microglia even in wild type brains. Such cells still exist in *Sall1* mutant brains. **E-H.** Double immunostaining of Iba1 and CD206 in the corpus callosum of wild type (E,F) and *Sall1* knockout (G,H) mice at P8. Many double-positive (arrowheads) cells exist in the corpus callosum of wild type brains. Such cells still exist in *Sall1* mutant brains with a density. Scale bars, 100μm.

### Morphology of developing microglia is altered in *Sall1* knockout mice

In addition to the pattern of gene expression, microglia also change from amoeboid to ramified morphology during development (Wu, Wen, Shieh, & Ling, 1992), whereas white matter microglia in the corpus callosum of early postnatal brain exhibit more amoeboid morphology than the microglia in the gray matter at the same stage (Li et al., 2019). Therefore, morphology of microglia is a good indicator of their temporal and spatial diversity. Already at E16.5, *Sall1*-mutant microglia in the gray matter showed less ramified patterns than those in wild type controls (Fig.6A,E). At P1, P8 and P21, *Sall1* mutant microglia in gray matter of the cerebral cortex continued to show reduced complexity of their processes, which left much more open space unoccupied by these processes compared with mild type microglia (Fig.7B-D, F-H). We quantified the changes in three-dimensional morphology of microglia by several parameters by using a MATLAB-based algorithm, 3DMorph (York, LeDue, Bernier, & MacVicar, 2018). We found that Iba1+ cells in *Sall1* cKO gray matter of the primary somatosensory cortex had a reduced number of branch points and reduced branch length, as well as the reduced ratio of territorial volume divided by the cell volume (“ramification index”) (Fig.7I,J,K). Microglia in the corpus callosum in P8 and P21 wild type cortex looked less ramified than those in the gray matter (Fig.7C,D,L,M). In *Sall1* cKO brains, these microglia appeared less dense and less ramified (Fig.7L-O). Quantification showed a reduced number of branch points and reduced ramification index at P8, reduced number of branch points at P21 (Fig.7P,Q). Thus, microglia in *Sall1* mutant brain fail to show gradual ramification in both gray matter and white matter. In summary, the persistently high ratio of CD206-expressing microglia at later stages of development and the change in morphology both in gray matter and white matter microglia together indicate that SALL1 is required for the normal progression of microglia development.

**Figure 7.**
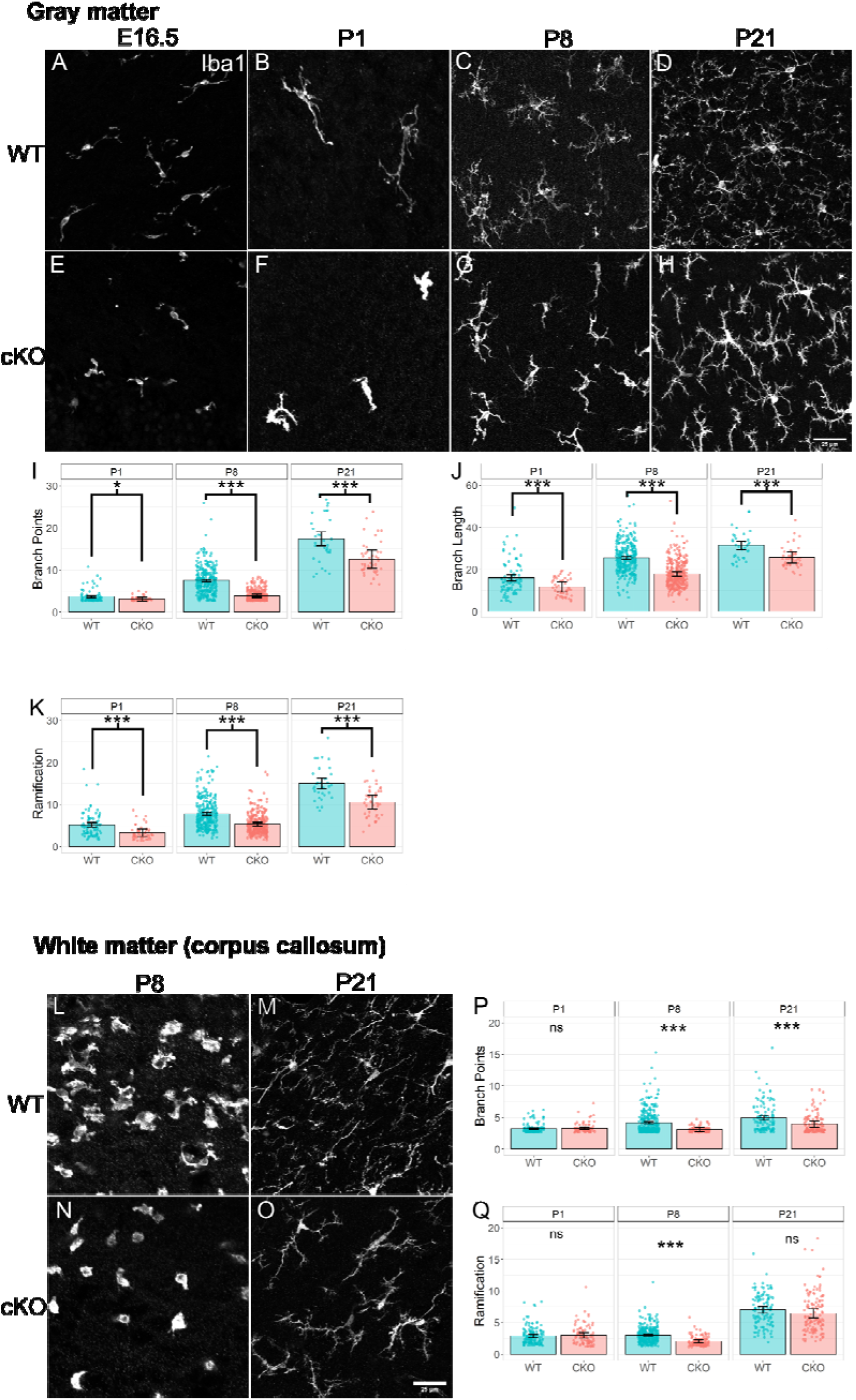
Altered microglia morphology in *Sall1* mutant brains. **A-H.** Immunostaining of Iba1 in the gray matter of the somatosensory cortex of wild type (A-D) and *Sall1* cKO (E-H) mice at E16.5 (A,E), P1(B,F), P8 (C,G) and P21 (D,H). *Sall1* mutant microglia show much less ramified morphology and coverage of space. All images were taken with a confocal microscope and represents a z-stack of 8 slices (0.49μm/slice). **I-K.** Quantitative analysis of microglia morphology using the 3DMorph program. Numbers of branch points (I), branch length (J) and ramification indexes (K) are compared between wild type (blue bars) and *Sall1* cKO (pink bars) primary somatosensory cortex. **L-O.** Immunostaining of Iba1 in the white matter of the somatosensory cortex of wild type (L,M) and *Sall1* cKO (N,O) mice at P8 (L,N) and P21 (M,O). *Sall1* mutant microglia show much less ramified morphology and coverage of space. All images were taken with a confocal microscope and represents a z-stack of 8 slices (0.49μm/slice). **P,Q.** Quantitative analysis of microglia morphology using the 3DMorph program. Numbers of branch points (P) and ramification indexes (Q) are compared between wild type (blue bars) and *Sall1* cKO (pink bars) primary somatosensory cortex.

### Sustained requirement of SALL1 in microglia development

When *Sall1* was deleted by the *Lyve1^Cre^* driver or by the *Cx3cr1^CreER^* driver with the CreER activation at E11.5, *Sall1* was removed in microglia from the beginning of its expression at E12.5. We next tested if SALL1 continues to be required for timely development of microglia at later stages. For this purpose, we delayed the deletion of *Sall1* gene by using the *Cx3cr1^CreER^* driver with the tamoxifen administration at P1 to activate the CreER protein. When we analyzed such brains at P8, a majority of Iba+ parenchymal cells expressed CD206 (Fig.S1E-H) similar to *Sall1* mutants that lacked SALL1 since E12.5 (Fig.4A-H). Morphology of microglia was also abnormal at P8 with a delayed deletion of *Sall1* at P1 (Fig.S1I,J). Thus, SALL1 continues to be required for normal progression of microglia maturation in early postnatal CNS.

## Discussion

In this study, we first found that the macrophage mannose receptor CD206 was expressed in early embryonic microglia, which was downregulated by E13.5. We then used *Lyve1^Cre^* or *Cx3cr1^CreER^* driver mice to delete the zinc finger transcription factor *Sall1* in order to test the intrinsic requirement of this gene in microglia development. With the *Lyve1^Cre^* driver, *Sall1* was deleted in microglia from the earliest stage of its expression at E12.5. In these mice, microglia showed a persistent expression of CD206 and reduced ramification in morphology into postnatal stages. In addition, delayed deletion of *Sall1* at a neonatal stage with the *Cx3cr1^CreER^* driver resulted in re-expression of CD206 and less ramified morphology compared with wild type controls.

A recent study (Utz et al., 2020) showed that in E10.5 and E12.5 brains, CD206 is expressed in 80-90% of *Cx3cr1*-expressing non-microglial macrophages in the meninges and choroid plexus, but only in 10% of parenchymal *Cx3cr1*-expressing cells. Since we used an anti-CD206 antibody similar to what the Utz et al. study used, the higher percentage of CD206 positivity in our study at these early stages suggests a higher sensitivity of immunostaining or lower thresholds to determine the positivity in our analysis. Although staining for CD206 in the brain parenchyma was weaker than in the meninges, we used two different threshold values in the quantitative analysis using the FIJI program and found clearly detectable expression with both thresholds. The dramatic reduction of the percentage of Iba1+ cells that also express CD206 between E12.5 and E13.5 in wild type brains is consistent with a RNA-sequencing study showing that *Mrc1* expression drops abruptly by 8 times between E12.5 and E13.5 in *Cx3cr1*+ cells in the brain (Matcovitch-Natan et al., 2016). In our analysis, all regions except the retina showed persistent expression of CD206 in *Sall1* mutant Iba1+ cells at E13.5 and later, whereas very few cells expressed CD206 in wild type brains at the same stage with the exception of some regions along axon tracts and around the blood vessels (perivascular macrophages) in postnatal animals. In addition, a majority of *Sall1* mutant Iba1+CD206+ cells also expressed microglia-specific markers P2RY12 and Tmem119 but not the BAM marker LYVE1, suggesting that these cells at least partially retained the identity as microglia but have an aberrant profile of gene expression consistent with immature microglia. This is further supported by the altered morphology of the mutant Iba1+ cells. Alteration of CD206 expression and microglia morphology was also observed at P8 when *Sall1* was deleted at P1, not during embryogenesis. This result shows that SALL1 is required not only during the early stage of microglia development, but is continuously required for maintaining the suppression of CD206 and morphological maturation during postnatal stages. Buttgereit et al. (2016) showed that deletion of *Sall1* in adult *Cx3cr1*+ cells resulted in altered morphology of microglia as well as their patterns of gene expression, including the upregulation of *Mrc1*. This finding together with ours collectively demonstrates that SALL1 plays critical role through the entire ontogeny of microglia into the adulthood. Further analysis of transcriptomic, epitranscriptomic and epigenetic status of these cells both at the single-cell level or with spatial genomics will provide detailed information as to how SALL1 regulates the key downstream events of microglia development. In particular, it will be important to determine if SALL1 directly inhibits transcription of *Mrc1* in developing microglia. Direct transcriptional targets of SALL1 in microglia have not been identified.

In our study, we assessed the persistent expression of CD206 in *Sall1*-deleted microglia in several regions in the CNS including the neocortex, lateral and medial ganglionic eminences, thalamus and the retina. Of these regions, the retina was an exception in that CD206 was downregulated significantly even in the absence of SALL1. A recent report has found reduced complexity of microglia morphology in *Sall1* mutant retina, but the morphology became qualitatively indistinguishable at postnatal stage (Koso et al., 2016; Koso, Nishinakamura, & Watanabe, 2018). This contrasts with our findings in the neocortex, where *Sall1* deficiency had a continuous impact on both CD206 expression and morphology into postnatal stages. It is possible that *Sall1* is less critical in the retina for microglia development because of the compensation by another Sall family transcription factor, *Sall3*.

This study has revealed an aspect of microglia diversity in developing CNS through the expression of *CD206/Mrc1*, and opens up further questions for future studies. First, a large subpopulation of Iba1+ cells in brain parenchyma expressed CD206 as early as at E10.5. Starting at E13.5, these CD206+Iba1+ cells were rarely found in the parenchyma except near axon tracts including the external capsule in embryos and corpus callosum in early postnatal brains. Whether these CD206+ microglia at early and late stages in CNS development share the same origin as CD206-negative cells has not been addressed. CD206, which is encoded by the *Mrc1* gene, is a transmembrane mannose receptor. Both *in vivo* and *in vitro* studies indicate that CD206 plays a role in inflammation and phagocytosis (Martinez-Pomares, 2012), although their roles in brain development have not been demonstrated. Thus, it is important to determine whether and how persistent expression of CD206 in *Sall1* mutant mice affects various aspects of neural development, including the viability of neural progenitor cells and neurons, as well as pruning of synapses. It has been shown that CD206+ white matter microglia in postnatal cortex also express the cell surface receptor protein,TREM2 (Chertoff, Shrivastava, Gonzalez, Acarin, & Giménez-Llort, 2013). This cell population potentially overlaps with recently described microglia subpopulations PAM (proliferative-region-associated-microglia) (Li et al., 2019) or ATM (axon tract-associated microglia) (Hammond et al., 2019), which might have specific functions in early postnatal brain development such as myelination (Wlodarczyk et al., 2017; Hagemeyer et al., 2017) and neuronal survival (Ueno et al., 2013) as well as neurogenesis and gliogenesis (Shigemoto-Mogami, Hoshikawa, Goldman, Sekino, & Sato, 2014). We detected changes in morphology of white matter microglia and altered distribution of CD206 in *Sall1* mutant mice, implying that SALL1 might regulate the above events during postnatal neural development.

We found that the *Lyve1^Cre^* driver (Pham et al., 2010) caused recombination not only in BAMs and but also in microglia from their earliest presence in the CNS. Recent reports (Kim et al., 2021; Utz et al., 2020) predicted that the *Lyve1^Cre^* driver would not cause recombination in microglia. However, (Lee et al., 2016) showed that the *Lyve1^Cre^* driver traces erythromyeloid progenitor cells in the yolk sac, which are considered to be the major source of microglia and BAMs (Ginhoux et al., 2010; Gomez Perdiguero et al., 2015; Kierdorf et al., 2013; Schulz et al., 2012). Thus, our results support these previous studies and add a genetic tool to manipulate microglia development *in vivo*.

Microglia play many roles in neural development from neurogenesis to cell survival, as well as synapse formation and elimination. Because of the dynamic changes in gene expression and morphology of microglia across time and space, it is expected that neural cells and microglia constantly interact with each other and modify each other’s developmental programs. In order to determine the underlying molecular mechanisms for such interactions, it is essential to independently manipulate each system in an intrinsic manner. We expect that the microgliaspecific *Sall1* mutant mice will be a useful platform to test the roles of normal microglial development in many aspects of neural cell development and to reveal the consequences of aberrant microglia development in brain functions and behavior later in life.

## Materials and Methods

### Mice

Care and experimentation on mice were done in accordance with the Institutional Animal Care and Use Committee at the University of Minnesota. Noon of the day on which the vaginal plug was found was counted as embryonic day 0.5 (E0.5), and the day of birth (E19.5) was designated as postnatal day 0 (P0). To generate mice with a conditional deletion of *Sall1*, *Lyve1*^*Cre/*+^;*Sall1*^*flox/*+^ mice were bred with *Sall1*^*flox/*+^ mice. *Sall1^flox/flox^* mice (Yuri et al., 2009; Kanda et al., 2014) were obtained from Yasushiko Kawakami (University of Minnesota). *Lyve1^Cre^* mice (Pham et al., 2010) were obtained from Jackson Laboratory. *Lyve1*^*Cre/*+^; *Sall1*^+/+^ mice were used as wildtype (WT) littermates and compared to *Lyve1*^*Cre/*+^; *Sall1^flox/flox^* as *Sall1* conditional knockout mice (cKO). Mice were kept in mixed background and embryos or pups of either sex were used. The *Lyve1^Cre^* allele contains *EGFP-IRES-Cre* construct in the 3’ untranslated region of the *Lyve1* gene, but in our experimental condition EGFP was only detectable by using an anti-EGFP antibody, and was not used in our analysis. We also produced and analyzed conditional *Sall1* mutant mice using *Cx3cr1^CreER^* (Yona et al., 2013) as the driver. In this case, *Cx3cr1*^*CreER/*+^; *Sall1*^+/+^ mice were used as wildtype (WT) littermates and compared to *Cx3cr1*^*CreER/*+^; *Sall1^flox/flox^* as *Sall1* conditional knockout mice (cKO). In some experiments, the tdTomato Cre reporter (Ai14) (Madisen et al., 2010) allele was included. When the *Cx3cr1^CreER^* was used, tamoxifen was administered either at E11.5 or at P1. For the E11.5 administration, pregnant dams were given an oral gavage of 3mg (0.6ml) tamoxifen dissolved in corn oil. For the P1 administration, individual pups were subcutaneously injected with 0.25mg (0.05ml) tamoxifen dissolved in corn oil by using a 30G needle.

### Tissue preparation

Embryos at or younger than E14.5 were removed from the dam and heads were immersion-fixed in 4% paraformaldehyde (PFA) (dissolved in 0.1M phosphate buffer (PB)) for 30 minutes. E16.5 mice were removed from the dam and intracardiacally perfused with 4% PFA, brains dissected in 0.1M PB, and immersion fixed in 4% PFA for 30 minutes. Postnatal mice were intracardiacally perfused by 4% PFA, brains dissected in 0.1M PB, and immersion fixed in 4% PFA for 60 minutes for P8 or 120 minutes for P14 and P21. P8 brains were first perfused by 0.1M PB before PFA. All brains were then submerged in 30% sucrose/ 0.1M PB overnight at 4°C. Brains younger than P14 were frozen in TissueTek OCT compound on dry ice. Frozen brains were stored at −80°C until usage. P14 and P21 brains were kept in 0.1M PB until 2 days before usage, and then submerged in 30% sucrose.

### Immunohistochemical staining

Immunohistochemical staining was performed based on (Vue et al., 2007). Embryonic brains were cut at 20μm thickness, and P1 and P8 brain at 40μm thickness, with a cryostat. P14 and P21 brains were cut with a sliding microtome at 50μm thickness. Sections on slides were left to dry on a slide warmer for 30 minutes prior to starting immunohistochemistry. For counting CD206+ cells, matched WT and cKO littermates were sectioned onto the same slides to reduce variability of immunohistochemistry between slides. Sections were then rinsed in 0.1M phosphate-buffered saline (PBS), immersed in boiled 10mM citrate buffer (pH 6) for 5 minutes prior to blocking with 3% donkey serum / 0.3% Triton X-100/ PBS for 1 hour. Primary antibodies were then added at appropriate dilutions and incubation was performed overnight at 4°C. The following primary antibodies were used:

| *Protein* | *Dilution* | *Manufacturer* | *Product #* | *Species* |
| --- | --- | --- | --- | --- |
| Iba1 | 200 | Wako Chemicals | 019-19741 | Rabbit |
| Iba1 | 200 | Synaptic Systems | 234004 | Guinea Pig |
| CD206 | 200 | BioLegend | 141702 | Rat |
| PU.1 | 50 | Santa Cruz | sc-390405 | Mouse |
| SALL1 | 200 | Abcam | ab31526 | Rabbit |
| P2RY12 | 100 | Anaspec | SQ-ANAB-77022 | Rabbit |
| Tmem119 | 200 | Abcam | Ab209664 | Rabbit |
| LYVE1 | 200 | R&D Systems | AF2105 | Goat |
| RFP | 200 | Rockland | 600-401-379 | Rabbit |
|  |  | Immunochemicals |  |  |
| tdTomato | 200 | Origene | AB8181 | Goat |

On the second day, after washing in PBS, sections were incubated with fluorochrome-conjugated secondary antibodies (Cy2, Cy3, or Cy5, from Jackson ImmunoResearch, West Grove, PA) for 1 hour, followed by DAPI counterstaining. After dehydration in ascending concentrations of ethanol and clearing in xylene, slides were mounted in DPX mounting medium (EM Sciences, Hatfield, PA).

### Imaging

For cell counting, images of sections that underwent immunohistochemistry were taken using an E800 upright microscope (Nikon, Tokyo, Japan) with a Retiga EXi camera (Qimaging, Burnaby, BC, Canada). Images were obtained with the Micromanager program (Edelstein et al., 2014) using the Multi-Dimensional Acquisition tool and saved as 12-bit OME-TIFF files. The exposure time for a given antibody was selected to achieve the maximum signal-to-noise ratio without reaching maximal intensity; this exposure time was identical for each slide with the same antibody. For morphology analysis, images were taken with a confocal microscope (see below under <u>Analysis of microglia morphology</u>).

### Binning

Images were entirely processed within the FIJI Images acquired with micro-manager were opened with FIJI (Schindelin et al., 2012) and the bin overlaid (FIJI_BinningMacro_MicroManager.ijm).

### Thresholding and cell counting

Images were processed with a custom ImageJ macro, which would batch-process all images with identical parameters across paired littermates. The process was 1) simple paraboloid background subtraction 2) normalization of signal intensity 3) thresholding by manually determined percentage 4) default watershed algorithm (FIJI_Thresholding-ITCN.ijm). Two threshold values (0.15 and 0.05) were used to count CD206+ cells that are also Iba1+.

### Statistical analysis for cell count data

Cell count data was compared between groups using a Welch’s two-sided t-test.

### Analysis of microglia morphology

Iba1-immunostained cortical sections were imaged with an Olympus FV1000 laser scanning confocal microscope with a 40x (1.3NA) objective. Cy3 was excited with a 543nm laser. Confocal stacks were taken at 1024×1024 pixels with the step size of 0.49μm. Images were processed with a custom ImageJ script and quantified with 3DMoprh (York et al., 2018) to measure various morphological characteristics including branch points and ramification index, a simplified measure of ramification that is calculated by measuring the territorial volume spanned by the microglia divided by the physical volume of the cell. Images were taken in the primary somatosensory and motor cortices.

## Acknowledgments

We thank Xiangyu Zou, Ellie Wheeler and Grace Heimdahl for technical assistance; Timothy Monko for help with image analysis; Marija Cvetanovic for critical reading of the manuscript; Yasuhiko Kawakami for providing *Sall1^flox/flox^* mice.

---

**Figure S1.**
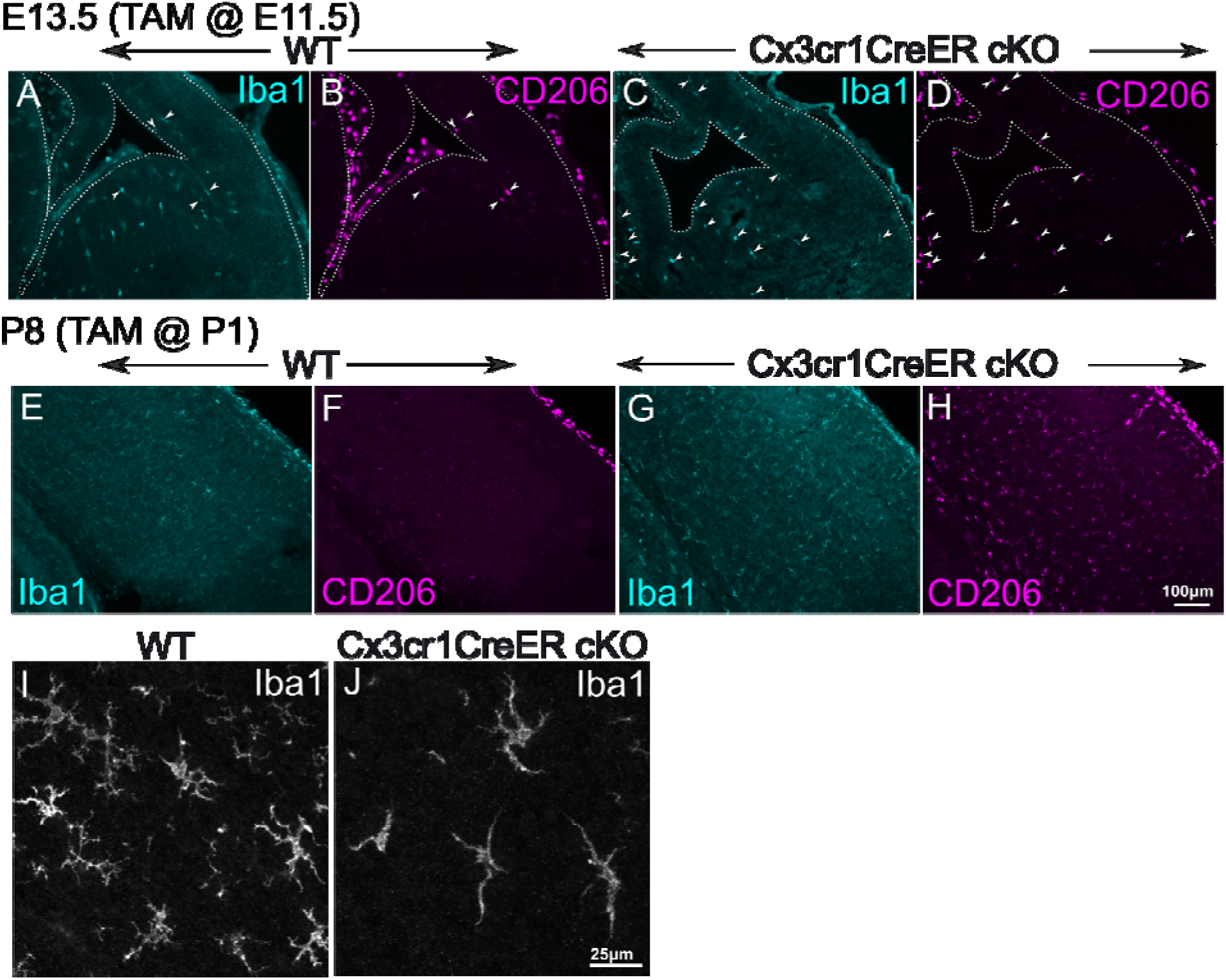
*Sall1* deletion with the Cx3cr1CreER driver. **A-D.** Double immunostaining of Iba1 and CD206 in wild type (A,B) and Cx3cr1CreER-mediated *Sall1* knockout (C,D) cortex at E13.5. Tamoxifen was administered at E11.5. Arrowheads indicate cells that express both Iba1 and CD206. **E-H.** Double immunostaining of Iba1 and CD206 in wild type (E,F) and Cx3cr1CreER-mediated *Sall1* knockout (G,H) cortex at P8. Tamoxifen was administered at P1. **I,J.** Iba1 immunostaining of wild type (I) and Cx3cr1CreER-mediated *Sall1* knockout (J) cortical gray matter at P8. Tamoxifen was administered at P1. Images were taken with a confocal microscope and represent a z-stack of 8 slices (0.49μm/slice).

